# Functionalization of lipid nanoemulsions with humanized antibodies using plug-and-play cholesterol anchor for targeting cancer cells

**DOI:** 10.1101/2025.06.29.662210

**Authors:** Valeria Jose Boide-Trujillo, Vincent Mittelheisser, Fei Liu, Olivier Lefebvre, Bohdan Andreiuk, Nicolas Anton, Jacky G. Goetz, Andrey S. Klymchenko

**Author notes:** **Correspondence and co-last authors:** Andrey S. Klymchenko, CNRS UMR_7021, Faculté de Pharmacie, Université de Strasbourg, Institut Thématique Interdisciplinaire SysChem, Illkirch F-67400, France., Jacky G. Goetz, INSERM UMR_S1109, Tumor Biomechanics Lab, Centre de Recherche en Biomédecine de Strasbourg (CRBS), Fédération de Médecine Translationnelle de Strasbourg (FMTS), 67000 Strasbourg, France. Web: www.goetzlab.fr. **Equal contribution:** VJBT, VM and FL have contributed equally to this work.

## Abstract

Lipid NEs are promising green nanocarriers for diagnostic and therapeutic applications, but their functionalization with biomolecules, such as antibodies, remains a challenge due to liquid nature of their core. Here, we developed an original plug-and-play strategy to graft an antibody (trastuzumab) at the surface of NEs, using components generally recognized as safe (GRAS). We synthesized a reactive derivative of cholesterol and a Biotin-PEG_3000_-Lysine linker, which can react within one-pot formulation to form an amphiphilic carbamate Biotin-PEG_3000_-Cholesterol. The cholesterol ensures anchorage of the linker, which effectively exposes biotin moiety at the surface of NEs for further antibody grafting using a biotin-neutravidin coupling. The reaction between the Biotin-PEG_3000_-Lysine linker and NPC-Chol was confirmed by ^1^H-NMR and absorption spectroscopy. The obtained biotinylated 50-nm NEs loaded with a near-infrared dye were successfully targeted to neutravidin-coated glass surfaces and imaged at the single-droplet level. The biotinylated NEs bearing the trastuzumab antibody targeted specifically HER2-amplified breast cancer models HCC-1954 and SKBR3, in contrast to control MDA-MB-231 (HER2-low) cells. Altogether, our study proposes an efficient methodology for grafting antibodies to the surface of NEs, which offers new opportunities of application of these green nanocarriers in biomedicine.

## Introduction

Cancer nanomedicine has initially emerged to improve the delivery selectivity of chemotherapeutics to reduce associated toxicities.^1, 2^ Among existing nanoparticles (NPs), nanoemulsions (NEs) constitute sustainable, biomimetic nanomaterials mainly composed of lipids and PEGs, which are materials *generally recognized as safe* (GRAS). ^3-5^ Their oil core serves as an ideal reservoir for encapsulating lipophilic compounds, including cytotoxic drugs and imaging agents.^6-8^ In particular, encapsulation of fluorescent dyes transforms them into bright fluorescence imaging agents.^6^ Dye-loaded NEs enable imaging vasculature and tumors,^8^ crossing brain blood barrier^9^ and intranasal brain delivery^10^ as well as tracking individual particles^11^ *in vivo*. Their long circulation time and potential of therapeutic molecules loading into their core render them particularly attractive for *in vivo* imaging and drug delivery.^6-8^ These nanocarriers are known to passively accumulate in the tumor through the enhanced permeability and retention (EPR) effect.^8, 12^ However, as defined in meta-analyses, this passive targeting remains suboptimal and thus requires implementing strategies for specific targeting.^13, 14^ Enhancing the targeting selectivity of drugs and contrast agents is a primary focus in cancer diagnostics and treatment to minimize systemic exposure and increase their efficacy.^15^ One major strategy has been the development of antibody-drug conjugates (ADCs), which merge the targeting specificity of monoclonal antibodies with the potency of cytotoxic drugs. These features have led to the widespread clinical adoption of ADCs, with routine applications for the treatment of hematological malignancies and HER2-amplified breast cancers due to their strong antitumoral activity.^16^ However, to maintain favorable pharmacokinetic and pharmacodynamic profiles, ADCs have relatively low drug/antibody ratios (≈ 4 molecules/antibody).^17^ To overcome this limitation, nanoparticles (NPs) delivery systems have been explored, which can carry >10 times as many small molecules as ADCs.^2, 18^ The same concerns contrast agents, where single NP can carry molecules, such as fluorescent dyes, this enhancing sensitivity of detection.^19, 20^ As such, combining the targeting selectivity of antibodies with the drug/contrast agent loading ability of NPs seems a logical step to advance this field. We recently confirmed the interest of developing such targeted formulations by highlighting that antibody-targeted nanoparticles demonstrate greater tumor uptake than non-decorated ones.^14^ However, functionalization of NEs with antibodies has been poorly explored to date.^6^

The major challenge in surface modifications of NEs is the dynamic nature of their liquid/liquid interface of NEs, which renders surface grafting poorly stable. In previous studies, NEs were functionalized with a peptide arginylglycylaspartic acid (RGD) using lipids^21^ or fatty acids,^22^ allowing the increased targeting of α_v_β_3_ integrin rich cells. However, this approach does not guarantee stable grafting for antibodies, because lipid/fatty acid residues may exchange with other lipid components of serum, as for instance was shown for other lipid nanoparticles.^23^ To overcome this limitation, we recently proposed to use NEs with amphiphilic polymers bearing biotin^24^ or an antibody.^25^ However, the limitation here is the use of non-biodegradable polymer, which raises concerns about long-term toxicity of this approach. Overall, antibodies are an excellent targeting ligand for delivery and imaging applications because of high affinity and specificity to the target.^26, 27^ They were found highly efficient for targeting different NPs in drug delivery applications.^14, 28, 29^ To achieve effective functionalization of NEs with antibodies and preserve GRAS characteristics and biomimetic nature, a “green” approach should be developed, which would involve exclusively natural/biocompatible components.

In the present work, we engineered antibody-targeted NEs to enhance tumor cell selectivity and improve their potential for clinical translation. To this end, we used biotinylation strategy which is commonly used in molecular biology and bio-nanotechnology for its rapid, efficient and robust bio-conjugation.^30, 31^ We synthesized a reactive derivative of cholesterol and a Biotin-PEG_3000_-Lysine linker, which can react within one-pot formulation to form an amphiphilic carbamate Biotin-PEG_3000_-Cholesterol. Cholesterol moiety was expected to ensure its firm anchorage in NEs, whereas the long hydrophilic PEG_3000_ chain was expected to expose biotin moiety at the surface of NEs for further antibody grafting using a biotin-neutravidin coupling. Successful NE biotinylation was confirmed by fluorescence staining on a glass surface. These biotinylated-NEs were further functionalized with biotinylated antibodies using neutravidin as a bridge. We selected trastuzumab as the targeting antibody due to its routine use as part of the therapeutic arsenal as a single agent or in ADCs formulations to treat HER2-amplified patients.^32^ Trastuzumab-functionalized NEs effectively recognized their cognate receptors and selectively accumulated into HER2-expressing cancer cell lines (in comparison to HER2-low ones). Overall, this study presents a highly biocompatible, modular and stable way to functionalize NEs surfaces with antibodies. These antibody-targeted NEs demonstrated enhanced and antigen-selective tumor cell targeting *in vitro*, suggesting that NEs are promising vehicles for targeted delivery and imaging.

## Results and discussion

### Design and synthesis of GRAS-compatible building blocks for NEs surface functionalization

We aimed at developing NEs decorated with an antibody to enable targeting tumor cell membrane expressing the cognate antigen. Given the liquid nature of the oil core, we considered that efficient anchoring of large biomolecules at the surface, can only be achieved by a linking with a highly hydrophobic anchor counterpart. To this end, we considered developing a plug-and-play appracoh, where a functional unit is “implanted” into the nano-droplet core by two cholesterol units though a long spacer. Cholesterol is a highly hydrophobic neutral lipid, which makes it attractive as a non-toxic anchor for NEs. We synthesized a reactive cholesterol derivative, (4-nitrophenyl) carbonate of cholesterol (NPC-Chol), which would allow attachment of the anchor group directly during preparation of NEs. We also designed and synthesized a PEG linker bearing lysine moiety on one end and biotin on the other. The two amino groups of lysine are expected to react with NPC-Chol thus anchoring the PEG linker in the oil core of the nano-droplets after their formulation. Due to sufficiently long PEG3000 linker, we expect that the biotin moiety would be exposed well outside of NEs surface. Then, biotinylated antibody is coupled to biotinylated-NEs though a neutravidin (avidin analogue) bridge. The selected antibody was trastuzumab, which is a commonly used therapeutic antibodies for targeting HER2 receptor

To address the current limitations of anti-HER2 ADCs and NPs, we aimed to develop trastuzumab-functionalized NEs capable of targeting their cognate antigen expressed on the surface of tumor cells. However, given the liquid nature of the NE oil core, efficient surface anchoring of large biomolecules such as the humanized IgG1 antibody trastuzumab requires conjugation to a highly hydrophobic anchor. To this end, we adopted a plug-and-play approach using GRAS compounds by designing reactive cholesterol derivative that can firmly anchor a functional linker within one-pot protocol. Cholesterol’s strong hydrophobic character and biocompatibility make it an attractive anchor for functionalization with an antibody at the NE surface. To this end, cholesterol was converted into reactive (4-nitrophenyl) carbonate of cholesterol (NPC-Chol) in one step by reaction with p-nitrophenyl chloroformate under basic conditions.^33^ Biotin-PEG_3000_-Lysine linker, which was synthesized in several steps (*see Supporting Information*) (**Figure 1**). Biotin was reacted with Mono-Boc-protected PEG_3000_ diamine to give compound **1**, which, after Boc removal in TFA, yielded compound **2**. Compound **2** was further reacted with Di-Boc-protected lysine to produce conjugate **3**. Then, the Boc groups of conjugate **3** were deprotected in TFA, yielding the final Biotin-PEG_3000_-Lysine linker (compound **4**). The identity of each synthesis intermediate as well as the final compound was confirmed by ^1^H-NMR and mass spectrometry (**Figure S1-5**). This linker is composed of three GRAS blocks. The lysine moiety, among the most common amino acids, are expected to react with NPC-Chol through their amino groups, thus anchoring the linker in the NE oil core after formulation. The spacer made of PEG, a widely used component in drug delivery, would allow exposure of derivatizable moieties away from the NE surface. The biotin, a water-soluble vitamin, is a convenient handle for further functionalization. Altogether, we designed and synthesized a GRAS, derivatizable anchor-linker system for subsequent conjugation with targeting ligands such as trastuzumab.

**Figure 1.**
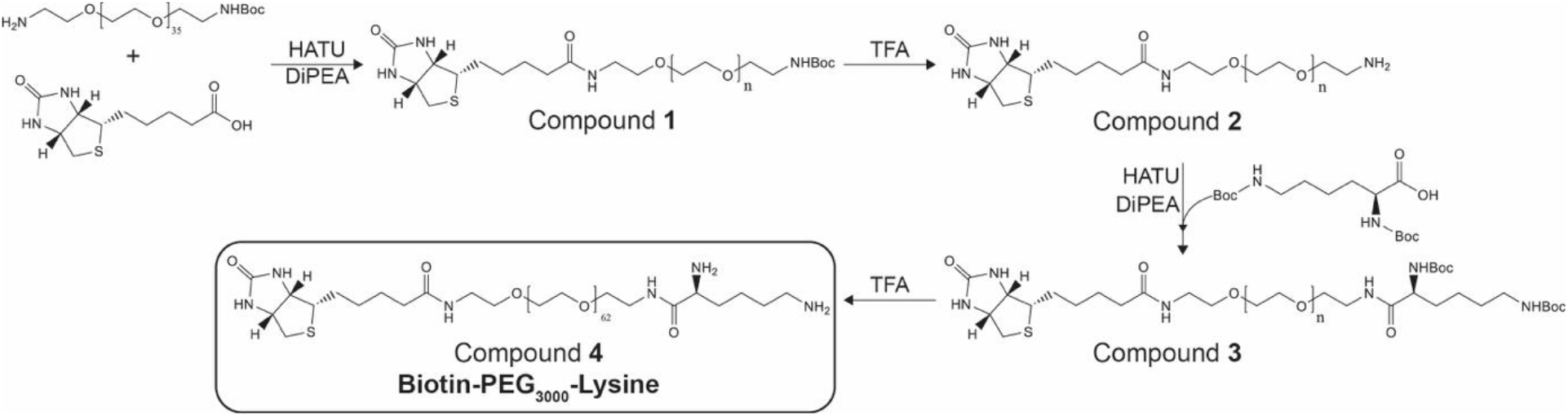
Synthesis of the Biotin-PEG_3000_-Lysine linker.

### Formulation of *in situ* biotinylated lipid nanoemulsion

In our approach, the reactive anchor NPC-Chol was first coupled with the linker Biotin-PEG_3000_-Lysine in THF in the presence of a base trimethylamine and Vitamin E acetate, which later on will serve as the oil core. After evaporation of the solvent and the base, the NMR spectra of the reaction mixture showed new peaks, corresponding to the liberated p-nitrophenol (**Figure 2B**). The integral of the new peaks suggested 18.5% conversion of the used NPC-Chol. Considering that complete reaction of one amino group of the linker with NPC-Chol would result in 23 % conversion of the latter, the observed 18.5% conversion suggests that at these conditions we grafted one cholesterol anchor per linker. Then, surfactant and water were added to the oil mixture containing the *in situ* generated Biotin-PEG_3000_-Chol, under intense stirring in order to obtain NEs by spontaneous nanoemulsification. In order to make independent verification that the NPC-Chol reacted with Biotin-PEG_3000_-Lysine, absorption spectrum of the obtained NEs mixture was recorded in order to identify the presence of released p-nitrophenol. Importantly, the reaction mixture showed the characteristic peak around 400 nm,^34^ whereas the control conditions without the linker showed negligible contribution of p-nitrophenol absorption. Thus, we confirmed that the reaction took place, which led to release of p-nitrophenol.

**Figure 2.**
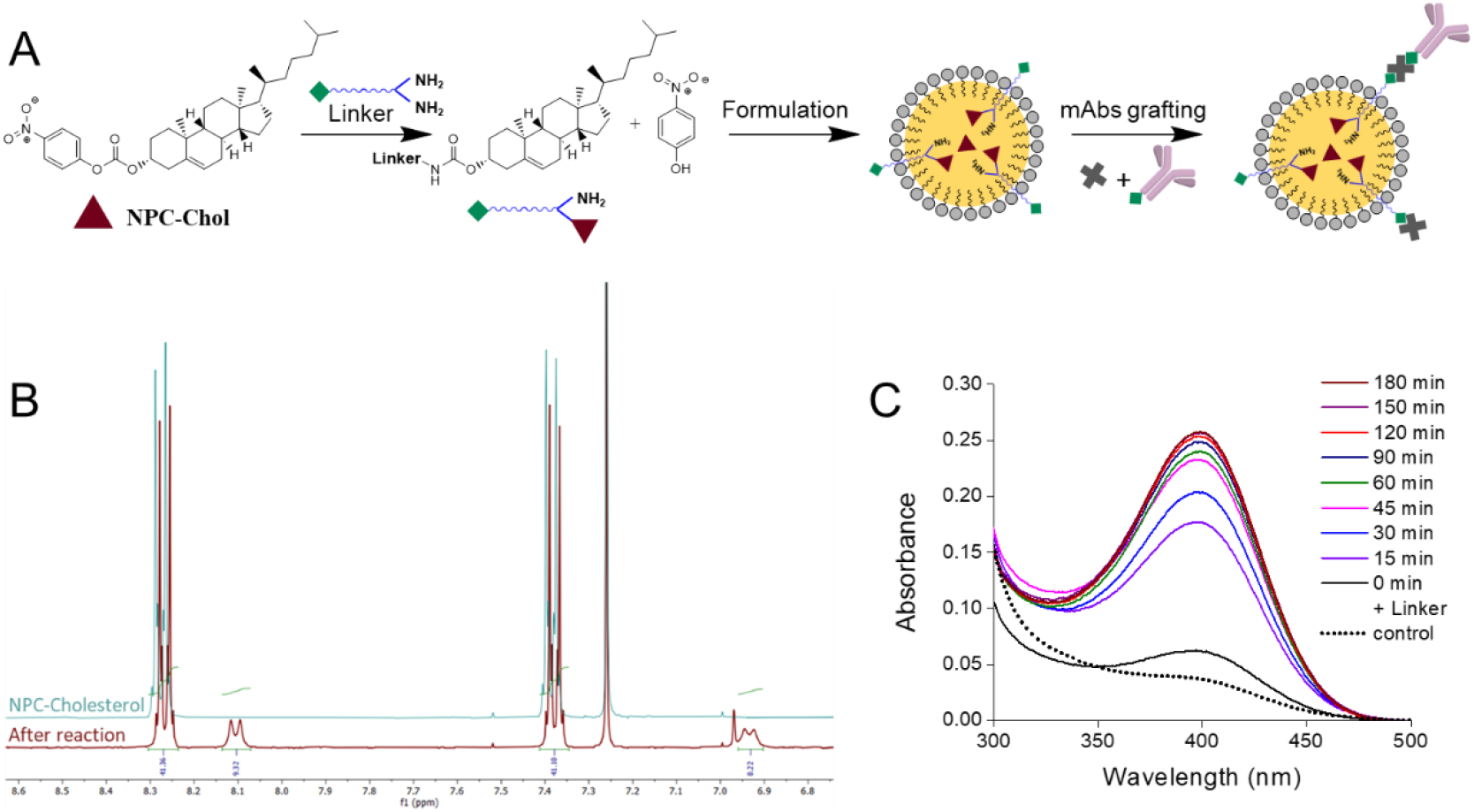
*In situ* biotinylation of lipid nanoemulsions. (**A**) *In situ* reaction between the Biotin-PEG_3000_-Lysine linker and the NPC-Chol generating the Biotin-PEG_3000_-Chol conjugate and releasing p-nitrophenol. (**B**) Superimposed ^1^H-NMR spectra of the NPC-Chol aromatic region, before and after the reaction with the Biotin-PEG_3000_-Lysine linker. (**C**) Absorption spectra of the released p-nitrophenol. The control experiment was done without Biotin-PEG_3000_-Lysine.

Then, to quench the non-reacted NPC-Chol, we incubated the NEs mixture with an excess of n-butylamine, which lead to further increase in the absorbance of p-nitrophenol. The reaction was incubated until no significant increase in absorbance was observed (**Figure 2C**). Assuming that after incubation with n-butylamine, 100% of NPC-Chol reacted with the amines liberating all p-nitrophenol, we could estimate that 17.1 % of NPC-Chol reacted Liotin-PEG_3000_-Lysine. This estimation is in line with the NMR data above, confirming formation of the linker-cholesterol conjugate. Then, the mixture was dialyzed to remove non-reacted linker, non-reacted butylamine and liberated p-nitrophenol. Dynamic light scattering (DLS) of the dialyzed product suggested the presence of nano-droplets of 64 ± 4 nm diameter with a polydispersity index of 0.2 ± 0.03 (Table S1). Altogether, we developed a one-pot green chemistry approach for NEs biotinylation using GRAS components, thereby reducing the risk of ligand detachment from the NEs surface.

### Validation of surface biotin activity via neutravidin binding assay

To assess the presence of active biotin units on the surface of NEs, we formulated biotinylated-NEs loaded with a hydrophobic cyanine 5.5 derivative and its bulky counterion tetraphenylborate (Cy5.5-LP/TPB) at 2 wt% in the oil phase (**Figure 3A**). This counterion has been shown to be essential to ensure efficient encapsulation of the dye within NEs with minimal dye leakage and aggregation-caused quenching.^8, 11^ According to DLS, the obtained Cy5.5-loaded biotinylated NEs were 53 ± 2 nm in diameter, with a polydispersity index of 0.16 ± 0.01 (**Table S1**). The absorption and emission spectra of these NEs showed single bands with maxima at 700 and 727 nm, were close to those for Cy5.5-LP in labrafac oil (**Figure 3B**). Using this Cy5.5-labeling, we characterized the biotinylated-NEs immobilized on glass surface at the single-particle level by fluorescence microscopy (**Figure 3B**). As a conjugation biomolecule for biotin, we used neutravidin, which is an avidin analogue presenting higher selectivity and lower non-specific binding.^35^ Hence, neutravidin is frequently used for immobilization of biotinylated molecules and nanoparticles on the surface.^24, 31, 36, 37^ To test interaction of neutravidin with biotin on NEs surface, we coated a glass surface with BSA-biotin, followed by a layer of neutravidin based on established protocols (**Figure 3A**).^36, 37^ With this surface at hand, we compared targeting specificity of biotinylated-NEs with control NEs without grafted biotin. Biotinylated-NEs were clearly visible on the glass surface by fluorescence microscopy in form of dots (**Figure 3D-E**). The density of dots and the overall intensity on the images increased with higher NEs concentration (**Figure 3D-E**). In case of control non-modified NEs, fluorescent dots were absent already for the highest used NEs concentration (**Figure 3C**). These results clearly show that the grafted biotin remains accessible and functionally active, as evidenced by its specific interaction with neutravidin, while non-specific interactions with the NEs with the surface are negligible.

**Figure 3.**
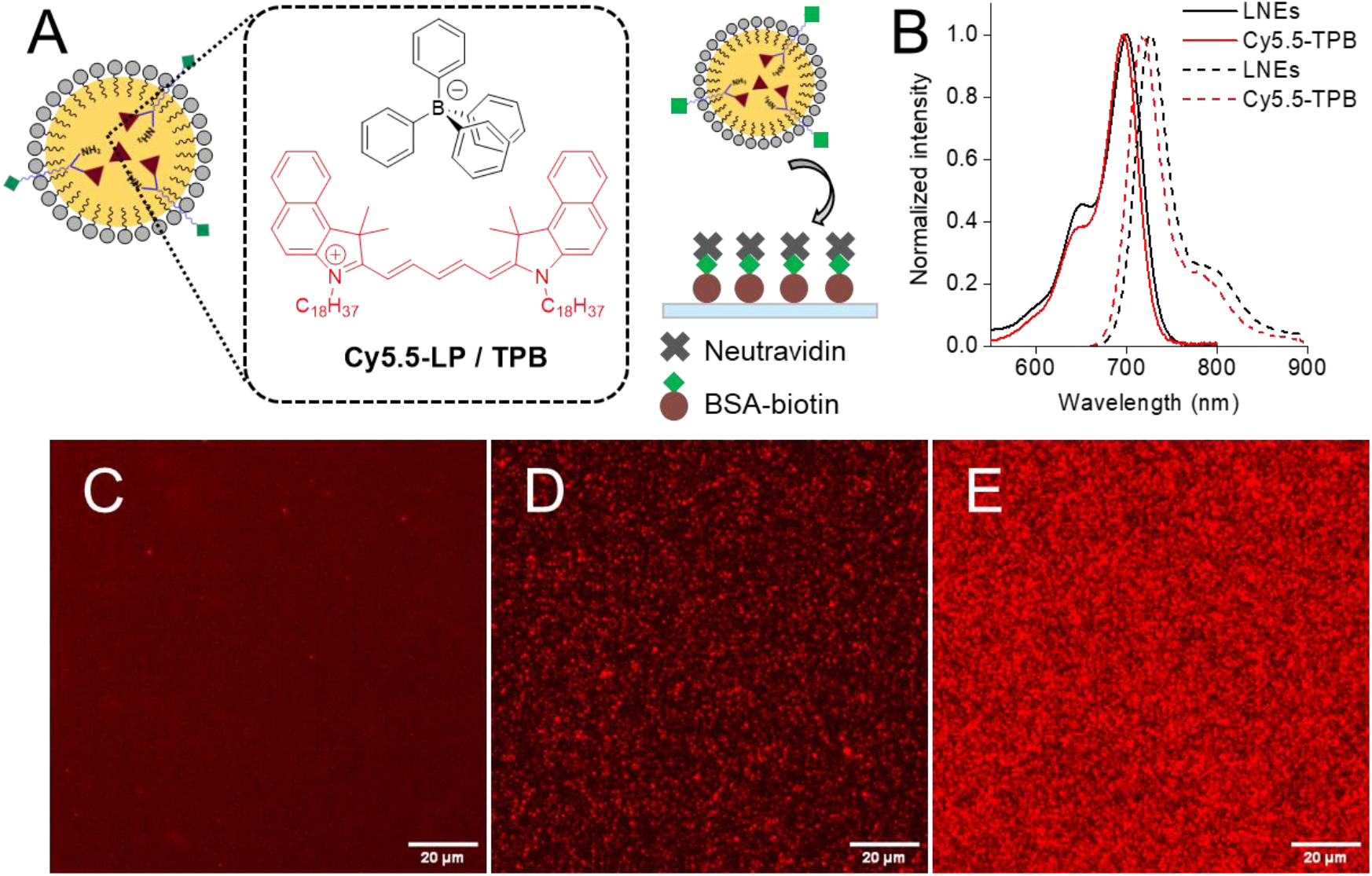
Validation of biotinylated nanoemulsions. (**A**) Schematic representation of Cy5.5-LP/TPB loaded NEs and their immobilization on a glass surface coated with BSA-biotin and neutravidin. (**B**) Absorption (solid line) and emission (dashed line) spectra of NEs loaded with Cy5.5-LP/TPB in water, and Cy5.5-LP/TPB in labrafac. (**C-E**) Fluorescence microscopy of neutravidin-coated glass surface incubated with bare NEs (30,000-fold dilution) (**C**) or biotin-decorated NEs (100,000-(**D**) or 30,000-fold (**E**) dilution). Scale bar = 20µm.

### Functionalization NEs with trastuzumab

Prompted by these results, we next generated NEs decorated with therapeutic antibodies to enable selective targeting of tumor cells expressing cognate antigens. For this purpose, trastuzumab – a clinically approved anti-HER2 antibody^34^ – was conjugated to biotin via its amino groups at physiological pH using a biotin-NHS ester,^34^ and unreacted reagent was removed by ultrafiltration. Efficient biotinylation was confirmed by strong membrane staining in a cell binding assay on HER2-amplified HCC1954, using Streptavidin-AF647 (SA-AF647) to detect accessible biotin sites (**Figure 4A**). The specificity of the biotin–streptavidin interaction was validated by pre-blocking with neutravidin, which abolished membrane-localized fluorescence (**Figure 4B**). Next, biotinylated trastuzumab was coupled to biotinylated NEs via a neutravidin bridge. This interaction is rapid, highly specific, and extremely stable (K_d_ ≈ 10^-15^ mol.L^-1^).^30, 31^ Unbound antibodies were then removed by size exclusion chromatography using Sephacryl S300 HR, which is specially adapted for separating relatively large macromolecules or globular proteins in the range 10 to 150 KDa. UV-visible absorbance spectra of the eluted fractions showed that trastuzumab-decorated NEs were recovered in early fractions (f3–f5), with coincident absorbance at 280 nm (protein) and 700 nm (Cy5.5-LP) (**Figure S6**). In contrast, free trastuzumab eluted in later fractions (f6–f8), showing absorbance only at 280 nm. This result suggests that the first fraction presenting both 280 and 700 nm absorbance (f3-f5) correspond to both NEs and conjugated trastuzumab, while the following fractions correspond to proteins (free antibodies and possibly neutravidin). The first fractions were combined and studied again by the same chromatography method. This time only a one band corresponding to the first fractions was observed, presenting absorbance in both 280 and 700 nm (**Figure S7**). Thus, size exclusion method allows separation of the of NEs from non-conjugated protein molecules. Importantly, antibody functionalization had negligible impact on NE hydrodynamic diameter (56 ± 2 nm) with preserved good polydispersity of 0.12 ± 0.03 (**Table S1**). To confirm that trastuzumab conjugation did not impair antigen recognition, we determined the apparent affinity constant (K_d_) of trastuzumab-decorated NEs by flow cytometry, using biotinylated NEs without antibody as a negative control. Fitting the binding data to a one-site Langmuir isotherm yielded an apparent K_d_ of 3.5 ± 1.8 nmol·L−^1^ (**Figure 4C**), consistent with previously reported values for trastuzumab in cell-based assays.^38, 39^

**Figure 4.**
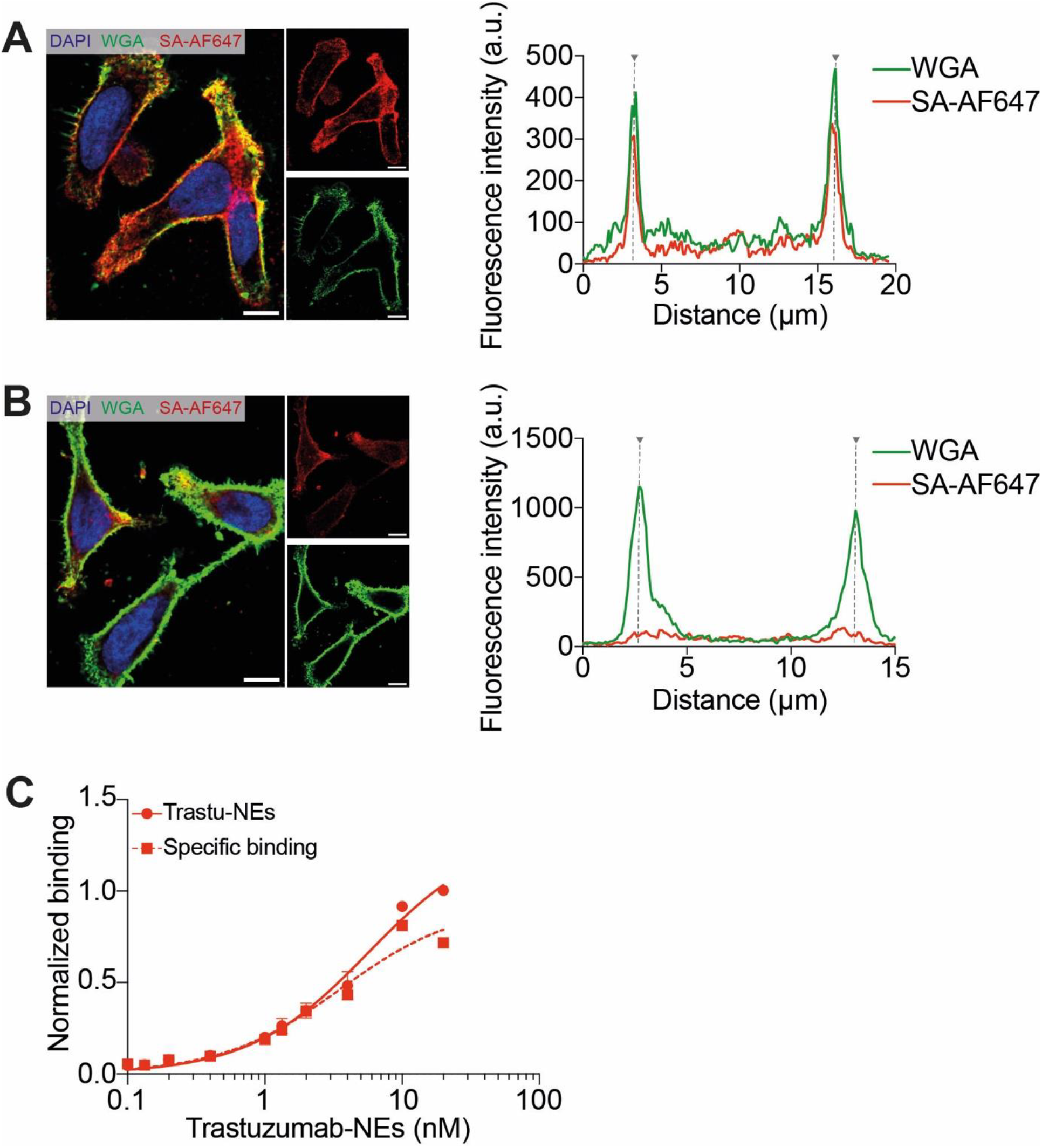
Validation of biotinylated trastuzumab on HER2-amplified HCC1954 cells. (**A-B**) Quantification of the trastuzumab-biotin membrane staining by following Streptavidin-AF647 (SA-AF647) fluorescence without (**A**) or with (**B**) pre-blocking with neutravidin. Cell membranes were labeled with Wheat Germ Agglutinin-AF488 (green). Scale bar is 10μm. Cell edges are marked by dot lines on the fluorescence profil plots. (**C**) Flow cytometry study determining the apparent binding affinity of the trastuzumab-functionalized nanoemulsions on HER2-amplified HCC-1954 cells.

### Trastuzumab-functionalized NEs target HER2-amplified tumor cells *in vitro* with high selectively

Building on these results, we next evaluated whether trastuzumab-functionalized NEs can target specifically tumor cells with high expression of HER2 receptor (HCC-1954) (**Figure 5A-B**) using confocal fluorescence imaging, of NEs loaded with Cy5.5-LP (2 wt% vs oil). Trastuzumab-functionalized NEs showed markedly increased internalization in HER2-amplified HCC-1954 cells compared to passive uptake by biotinylated NEs alone (**Figure 5C**). To confirm HER2 dependency, we assessed internalization in HER2^low^ MDA-MB-231 cells, where no detectable differences were observed between targeted and untargeted NEs (**Figure 5C**). To further validate the targeting selectivity, we extended the analysis to an additional HER2-positive cell line, SKBR3, which also showed preferential uptake of trastuzumab-targeted NEs over control formulations (**Figure S8**). To evaluate the biocompatibility of the formulation approach, we assessed whether NEs uptake induced cytotoxic effects. Treating both HER2-amplified and HER2^low^ cell lines with either trastuzumab-targeted or untargeted NEs had no effect on cell viability (**Figure S9**). Altogether, these results demonstrate the enhanced targeting selectivity of trastuzumab-functionalized NEs toward HER2-expressing tumor cells. This strategy provides a versatile and biocompatible platform for antibody-based surface functionalization of NEs to improve tumor-targeting efficiency.

**Figure 5.**
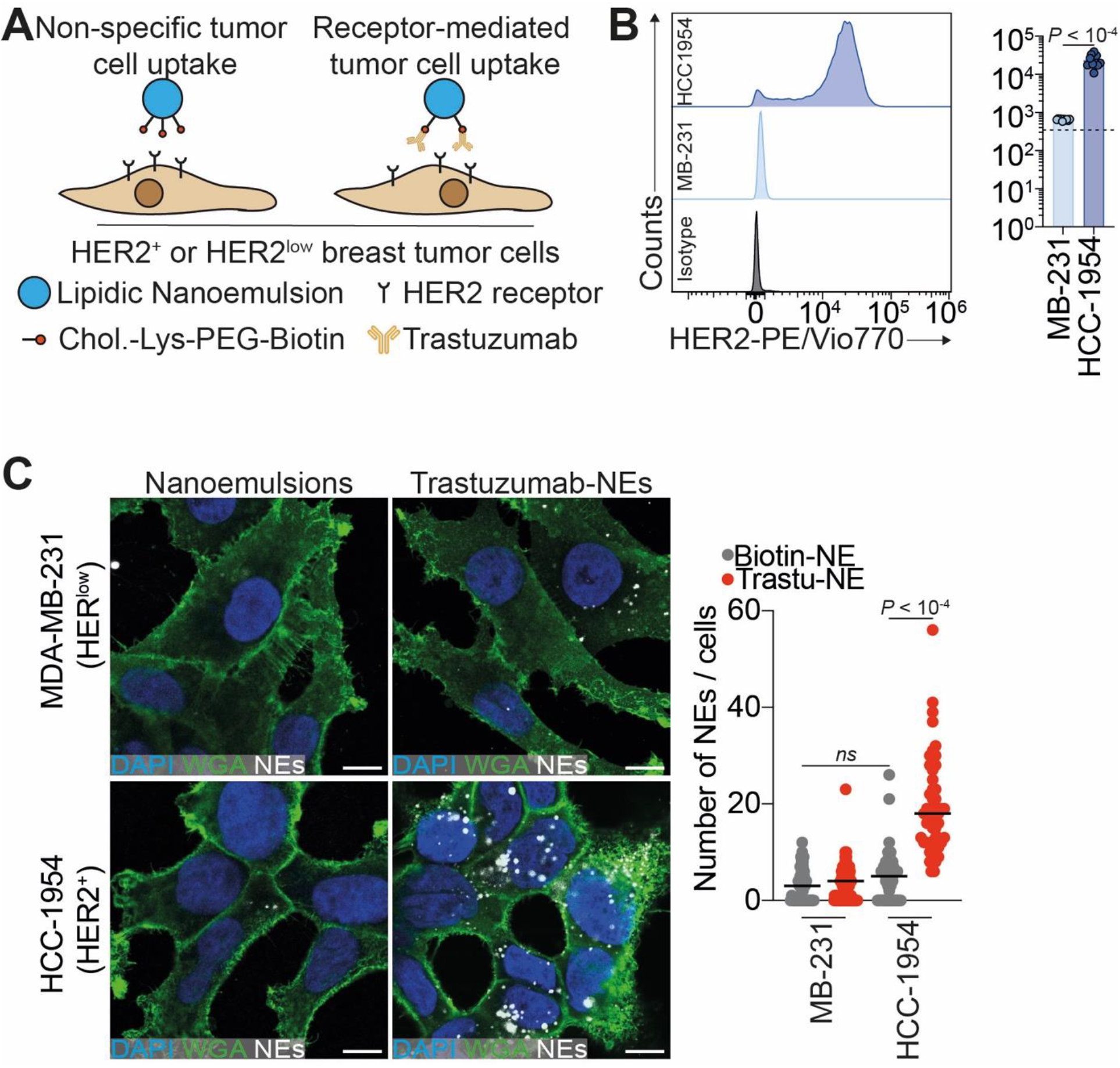
Trastuzumab-functionalized nanoemulsions target breast cancer cells in an antigen-antibody manner. (A) Biotinylated-NEs or Trastuzumab-targeted NEs interactions were assessed using HER2-amplified HCC-1954 cells or HER2^low^ MDA-MB-231 cells. (B) Flow cytometry analysis of HCC-1954 and MDA-MB-231 HER2 status. (C-D) Quantification of the nanoemulsion uptake in HER2^low^ (upper panel) and HER2-amplified (lower panel) cells. Cells incubated for 3h with 1.1 nM of Cy5.5-loaded (red) biotinylated NEs (right) or Trastuzumab-targeted NEs (left). Cell membranes were labeled with Wheat Germ Agglutinin-AF488 (green). Scale bar is 10 μm.

## Conclusions

Functionalization of lipid NEs with biomolecules for selective targeting remains a challenge. Here, we developed an original plug-and-play strategy to graft an antibody (trastuzumab) at the surface of NEs, using GRAS components. For this purpose, we synthesized a reactive derivative of cholesterol, cholesteryl (4-nitrophenol) carbonate (NPC-Chol), which can be readily encapsulated in the oil core of NEs and react with amines. We also designed a Biotin-PEG_3000_-Lysine linker with two amino groups of lysine at one end for reaction with NPC-Chol to form the amphiphilic carbamate Biotin-PEG_3000_-Chol. Cholesterol moiety was expected to ensure its firm anchorage in NEs. On the other end of the linker, the long hydrophilic PEG_3000_ chain is expected to expose biotin moiety at the surface of NEs for further antibody grafting using a biotin-neutravidin coupling. The reaction between the Biotin-PEG_3000_-Lysine linker and NPC-Chol was confirmed by ^1^H-NMR and absorption spectroscopy. Using one-pot approach, the obtained conjugate with oil and surfactant were formulated into 50-nm NEs, which were expected to bear biotin ligand. The successful biotinylation was confirmed by fluorescence microscopy: biotinylated-NEs loaded with a near-infrared dye were successfully targeted to neutravidin-coated glass surfaces in contrast to non-modified NEs. The obtained fluorescent biotinylated-NEs were grafted with biotinylated anti-HER2 (trastuzumab) monoclonal antibodies via a neutravidin bridge. The described approach presents several unique features compared to previously proposed methods. Compared to the previously proposed method based on amphiphilic polymers^24, 25^ the present approach uses GRAS and biodegradable components, which addresses the issues of safely. On the other hand, compared to methods based on fatty acids ^22^ or PEGylated lipids,^21^ the use of cholesterol increases the strength of anchoring with lower chance of exchange in biological medium due to higher hydrophobicity of the latter.

The targeting selectivity of the trastuzumab-decorated NEs was evaluated *in vitro* by confocal microscopy on relevant breast cancer models HCC-1954 and SKBR3 (HER2-amplified) and MDA-MB-231 (HER2^low^). Targeted NEs presented higher cellular internalization compared to control NEs in corresponding positive cell lines. In contrast, in control cell lines with low HER2 expression, no internalization was observed for both targeted and non-targeted formulations. These results show receptor dependent targeting of our NEs functionalized with antibodies.

Altogether, our study showed that combining the high encapsulation properties of the NEs with the trastuzumab surface decoration allowed receptor selective targeting of HER2-amplified breast cancer cells. These easily tunable functionalized NEs systems provide a platform for conjugation of different monoclonal antibodies and encapsulation of a large variety of contrast agents and drugs and thus can be used for imaging, therapeutic and targeted drug delivery applications. However, further improvements are still required. Indeed, the streptavidin analogue neutravidin used to bridge antibodies to NEs can be immunogenic^40^ and could eventually be replaced by direct biorthogonal conjugation methods such as click chemistry.^25, 41, 42^ In this case, the same green chemistry could be applied, where the biotin unit is replaced with corresponding reactive group. In addition, the NEs used in this study are lipidic nanoparticles and thus should, according to our meta-analysis results, be targeted with fragments rather than full antibodies for significantly higher accumulation.^14^ These improvements will be further complemented with *in vivo* experiments of mice bearing tumor models in order to evaluate the full potential of functionalized NEs in biomedical applications.

## Supporting information

Supplemental material for publication

## Acknowledgements

We thank all members of the JGG and ASK labs for helpful discussions. ASK team and VJBT were supported by ANR SenEmul and by the Interdisciplinary Thematic Institute SysChem, via the IdEx Unistra (ANR-10-IDEX-0002), the CSC Graduate School (CSC-IGS ANR-17-EURE-0016) within the French Investments for the Future Program. FL was supported by China Scholarship council (CSC). Work and people in JGG lab are mostly supported by the INCa (Institut National Du Cancer, French National Cancer Institute), La Ligue contre le Cancer, the Association pour la Recherche contre le Cancer (ARC), the Fondation pour la Recherche Médicale (FRM), the National Plan Cancer initiative, the Region Grand Est, INSERM and the University of Strasbourg as well as from local donators (Rohan Athlétisme Saverne, Trailers De La Rose, Club Féminin Lampertheim). VM was funded by a Ph.D. fellowship from the French Ministry of Science (MESRI) and by the Foundation ARC. We thank Pascal Kessler and Ignacio Busnelli from PICSTRA (CRBS). We thank Alexandre Detappe for providing Trastuzumab.

## Table of contents entry

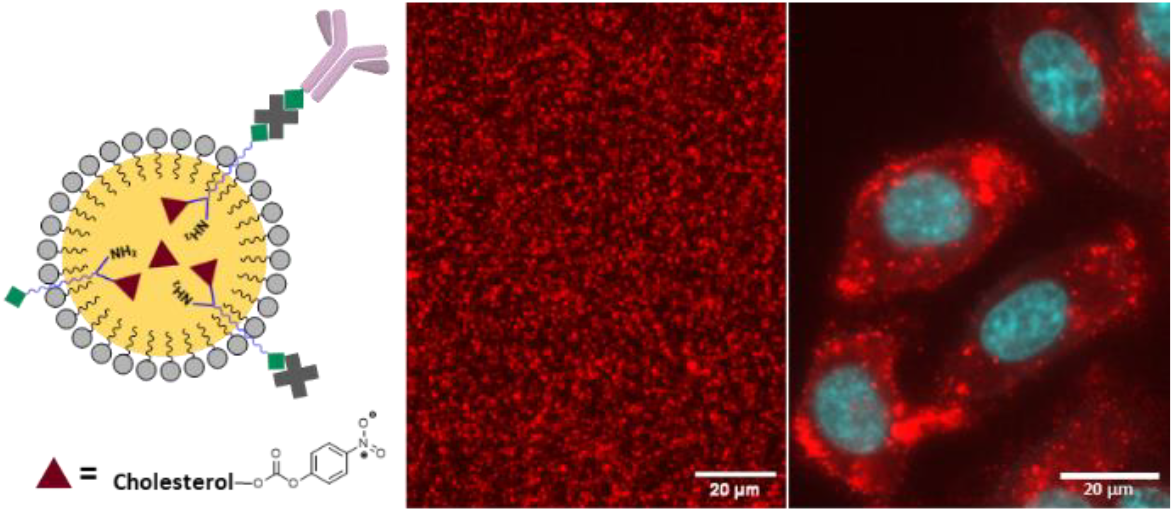

A plug-and-play functionalization of nanoemulsions with antibodies based on reactive cholesterol and bifunctional linker was proposed for specific cell targeting.

