## Supplemental material for publication for "Functionalization of lipid nanoemulsions with humanized antibodies using plug-and-play cholesterol anchor for targeting cancer cells"

### Materials and methods

#### Materials

Boc-NH-PEG-$NH_2$ (3 kDa) was purchased from Iris Biotech GmbH. Biotin (97%) and diBoc-lysine (98%) were acquired from abcr GmbH. 2-(7-Aza-1H-benzotriazole-1-yl)-1,1,3,3-tetramethyluronium hexafluorophosphate (HATU) was provided by Fluorochem. N,N-Diisoprozlethylamine (DiPEA) (>99%), Kolliphor® ELP, triethylamine, butylamine (99.5%), and BSA-biotin were supplied by Sigma-Aldrich. Trifluoro acetic acid (TFA) (99%), NHS-Biotin, and NeutrAvidin were obtained from ThermoFisher. Herceptin® (Trastuzumab) was purchased from the ICANS Pharmacy. Vitamin E acetate (>97.0%) was acquired from TCI. Dulbecco's Phosphate-buffer saline (DPBS) (without $Ca^{2+}$ and $Mg^{2+}$) was provided by Dominique Dutscher. Buffer preparations and formulation of lipid nanoemulsions (NEs) were done with filter-sterilized Milli-Q water. Borate buffer (pH 8.4) was prepared with sodium tetraborate (25 mM), NaCl (25 mM), and EDTA (1 mM). Phosphate buffer (pH 8.5) was prepared with dipotassium hydrogen phosphate (40 mM) and NaCl (150 mM). Final pH values were adjusted with 1 M hydrochloric acid or 1 M sodium hydroxide.

#### Equipment

Size distributions of NEs were determined by dynamic light scattering using a Zetasizer Nano series DTS 1060 (Malvern Instruments). Absorption Spectra were recorded on a Cary 500-UV-vis-NIR spectrophotometer (Agilent). NMR spectra were recorded at 20 °C on a Bruker Avance III 400 MHz NMR spectrometer (Bruker). Conjugation experiments were carried out in DNA low-binding tubes 1.5 mL (Eppendorf) at atmospheric pressure at room temperature unless otherwise stated. Ultrafiltration was carried out in centrifugal filters Amicon® Ultra 0.5 mL (Merck Millipore) with a molecular weight cut-off (MWCO) of 50 kDa. Centrifugation was carried out on a MiniSpin plus (Eppendorf) operating at 10.000 rpm at room temperature. Size exclusion chromatography was performed on an ÄKTA start (Cytivia) operating at a flow speed of 0.5 mL/min. Sephacryl S-300 High-Resolution (GE Healthcare) resin was used to pack the column. Epifluorescence microscopy images were recorded on an ECLIPSE Ti (Nikon). Confocal microscopy was performed on an inverted OIympus Spinning-disk.

### Synthesis

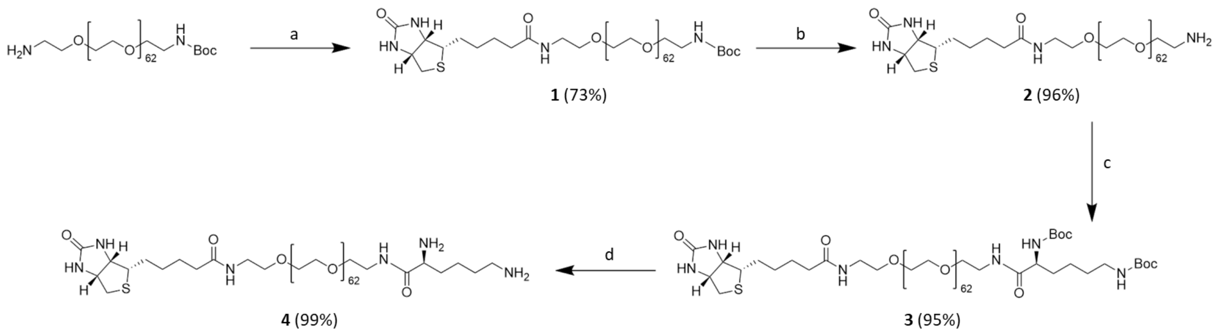

**Scheme 1.** Biotin-PEG-lysine synthesis (**4**): a) Biotin, HATU, DiPEA, DMF, 40 °C, overnight; b) TFA, DCM, rt, 3 hours; c) Boc-lysine(Boc)-OH, HATU, DiPEA, DMF, 40 °C, overnight; b) TFA, DCM, rt, 3 hours.

**Compound 1** - Biotin (5 eq., 62.3 mg., 0.25 mmol) and HATU (5 eq., 100.0 mg, 0.25 mmol) were dissolved in 5 mL of dry DMF. DiPEA (18 eq., 145 µL, 0.85 mmol) was added and stirred for 15 min. A 5 mL solution of Boc-H-PEG-$NH_2$ (1 eq., 140.0 mg, 0.03 mmol) in dry DMF was added to the previous mixture, and the final mixture was stirred overnight at 40 °C. The crude was purified by size exclusion chromatography LH-20 (DCM:MeOH, 1:1) to afford compound **1** (107.0 mg, 0.03 mmol, 73 %) as a colorless wax. $^1$H-NMR (400 MHz, $CD_3OD$) δ 4.50 (m, 1H), 4.31 (dd, J = 4.4, 7.8 Hz, 1H), 3.81 (t, J = 4.8 Hz, 2H), 3.70-3.58 (m, ~285H), 3.55 (m, 2H), 3.51 (t, J = 5.7 Hz, 1H), 3.46 (t, J = 4.7 Hz, 1H), 3.36 (m, 2H), 3.22 (p, J = 5.7 Hz, 3H), 2.93 (dd, J = 5.0, 12.7 Hz, 1H), 2.71 (d, J = 12.7 Hz, 1H), 2.23 (t, J = 7.4 Hz, 2H), 1.81-1.54 (bs, 4H), 1.48-1.41 (10H).

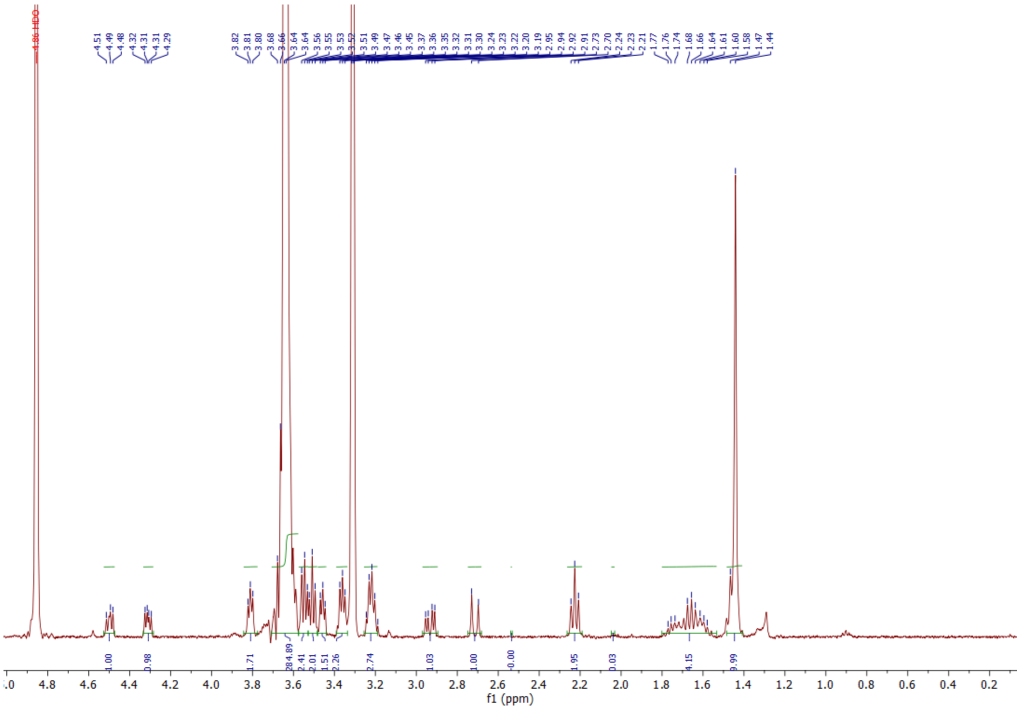


**Figure S1.** $^1$H-NMR of compound **1** in $CD_3OD$.

**Compound 2** - Compound **1** (1 eq., 107.0 mg, 0.03 mmol) was dissolved in dry DCM (2 mL). Trifluoroacetic acid (150 eq., 0.34 mL, 4.48 mmol) was added, followed by 3 hours of stirring at room temperature. DCM was removed under reduced pressure. Co-evaporation with methanol (1 mL) was done several times to remove the remaining trifluoroacetic acid, affording compound **2** (102.0 mg, 0.03 mmol, 96 %) as light-yellow wax. $^1$H-NMR (400 MHz, $CD_3OD$) δ 4.50 (dd, J = 4.9, 7.9 Hz, 1H), 4.31 (dd, J = 4.5, 7.9 Hz, 1H), 3.74-3.35 (m, ~278H),3.83-3.75 (m, 4H), 3.54 (t, J = 5.5 Hz, 2H), 3.46 (m, 1H), 3.36 (m, 3H), 3.20 (m, 3H), 2.93 (dd, J = 5.0, 12.8 Hz, 1H), 2.71 (d, J = 12.7 Hz, 1H), 2.23 (t, J = 7.3 Hz, 2H), 1.77- 1.53 (bs, 4H), 1.46 (q, J = 7.5 Hz, 2H).

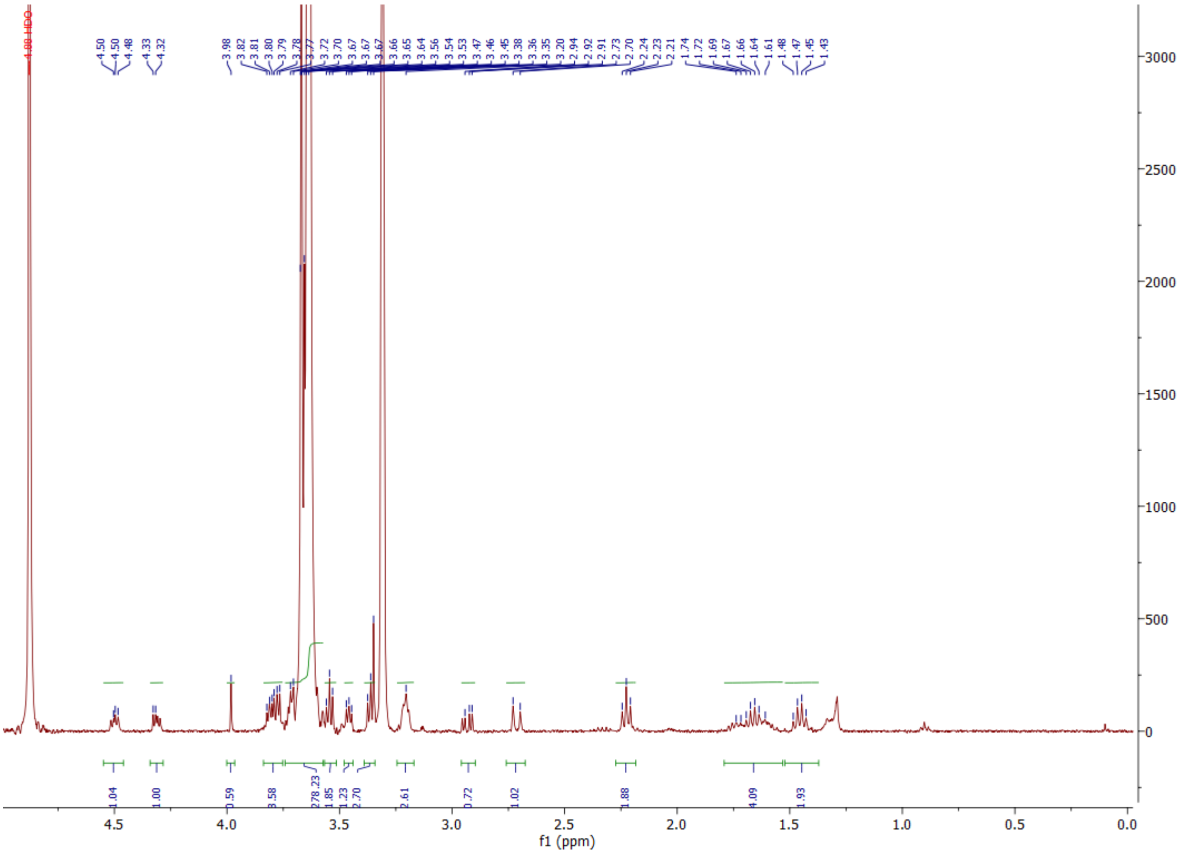


**Figure S2.** $^1$H-NMR of compound **2** in $CD_3OD$.

**Compound 3** - diBoc-lysine (3 eq., 51.4 mg., 0.09 mmol) and HATU (2.5 eq., 31.0 mg, 0.07 mmol) were dissolved in 5 mL of dry DMF. DiPEA (10 eq., 55 µL, 0.32 mmol) was added and stirred for 15 min. A 5 mL solution of compound **2** (1 eq., 102.0 mg, 0.03 mmol) in dry DMF was added to the previous mixture, and the final mixture was stirred overnight at 40 °C. The crude was purified by size exclusion chromatography LH-20 (DCM:MeOH, 1:1) to afford compound **3** (105.0 mg, 0.03 mmol, 95 %) as a colorless wax. $^1$H-NMR (400 MHz, $CD_3OD$) δ 4.50 (dd, J =4.8, 7.7 Hz, 1H), 4.31 (dd, J = 4.4, 7.9 Hz, 1H), 3.99 (m, 1H), 3.81 (m, 2H) 3.70-3.58 (m, ~315 H), 3.54 (t, J = 5.4 Hz, 5H), 3.45 (m, 2H), 3.37 (q, J = 5.5 Hz, 4H), 3.21 (m, 1H), 3.03 (t, J = 5.9 Hz, 2H), 2.93 (dd, J = 5.0, 12.7 Hz, 1H), 2.72 (d, J = 12.7 Hz 1H), 2.23 (t, J = 7.4 Hz, 2H), 1.81-1.54 (m, 7H), 1.52-1.40 (m, 25H).

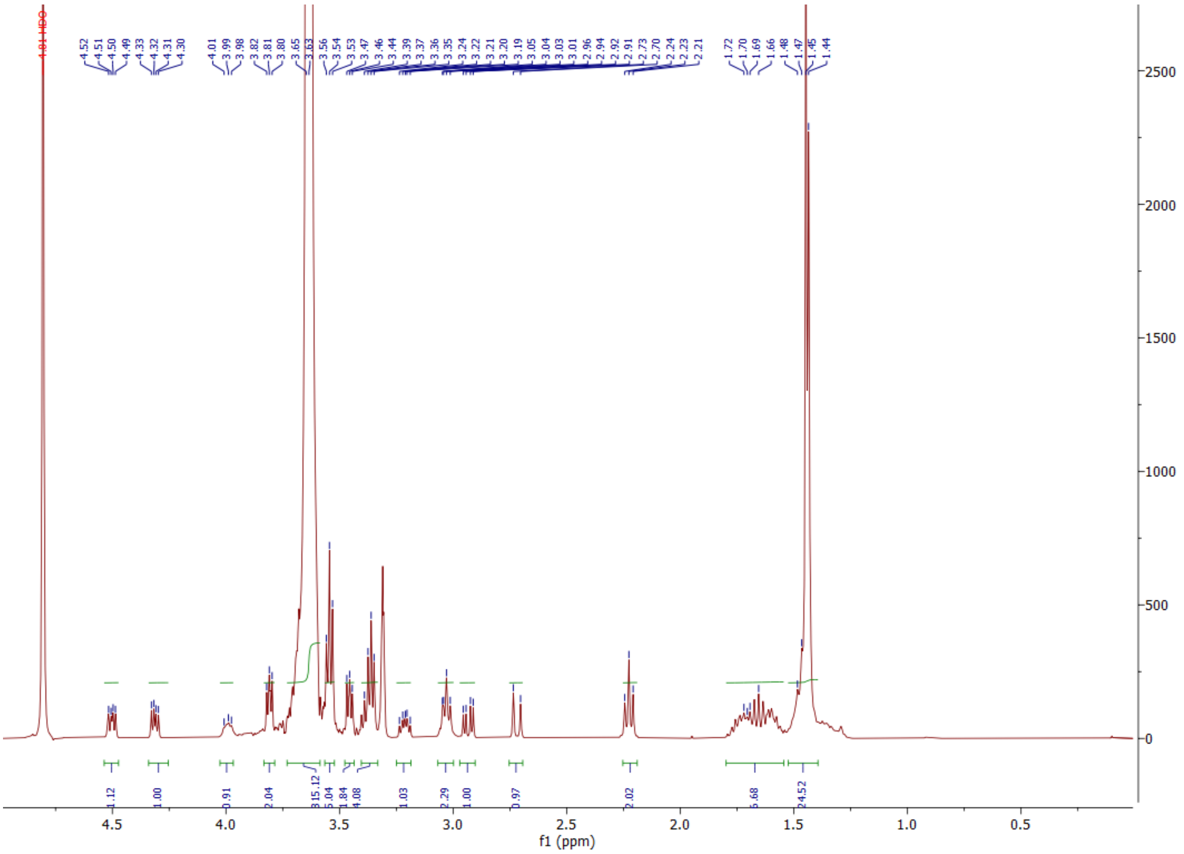


**Figure S3.** $^{1}$H-NMR of compound **3** in $CD_3OD$.

**Compound 4** - Compound **3** (1 eq., 105.0 mg, 0.03 mmol) was dissolved in 2 mL of dry DCM. Trifluoroacetic acid (150 eq., 0.34 mL, 4.48 mmol) was added, followed by 3 hours of stirring at room temperature. DCM was removed under reduced pressure. Co-evaporation with methanol (1mL) was done several times to remove the remaining trifluoroacetic acid, affording compound **4** (104.0 mg, 0.03 mmol, 99 %) as light-yellow wax. $^{1}$H-NMR (400 MHz, $CD_3OD$) δ 4.49 (dd, J = 4.8, 8.0, 1H), 4.30 (dd, J = 4.3, 7.8, 1H), 3.95 (t, J = 6.6 Hz, 1H), 3.8 (t, J = 4.8 Hz, 2H) 3.71– 3.56 (m, ~306 H), 3.54 (t, J = 5.4 Hz, 3H), 3.45 (m, 2H), 3.35 (m, 2H), 3.20 (dt, J = 5.2, 9.9 Hz, 1H), 3.02 (t, J = 7.9 Hz, 2H), 2.92 (dd, J = 5.0, 12.8 Hz, 1H), 2.70 (d, J = 12.4 Hz, 1H), 2.22 (t, J = 7.3 Hz, 2H), 1.91 (q, J = 7.9 Hz, 2H), 1.80-1.55 (m, 6H), 1.55 -1.40 (m, 4H). Mass data: $C_{144}H_{286}N_6O_{66}S$ (Exact mass expected 3187.92). Mass detected: 1617.70 and 1086.09 (calc.: $(M+2Na^+)/2$=1616.96 and $(M+3Na^+)/3$=1085.64).

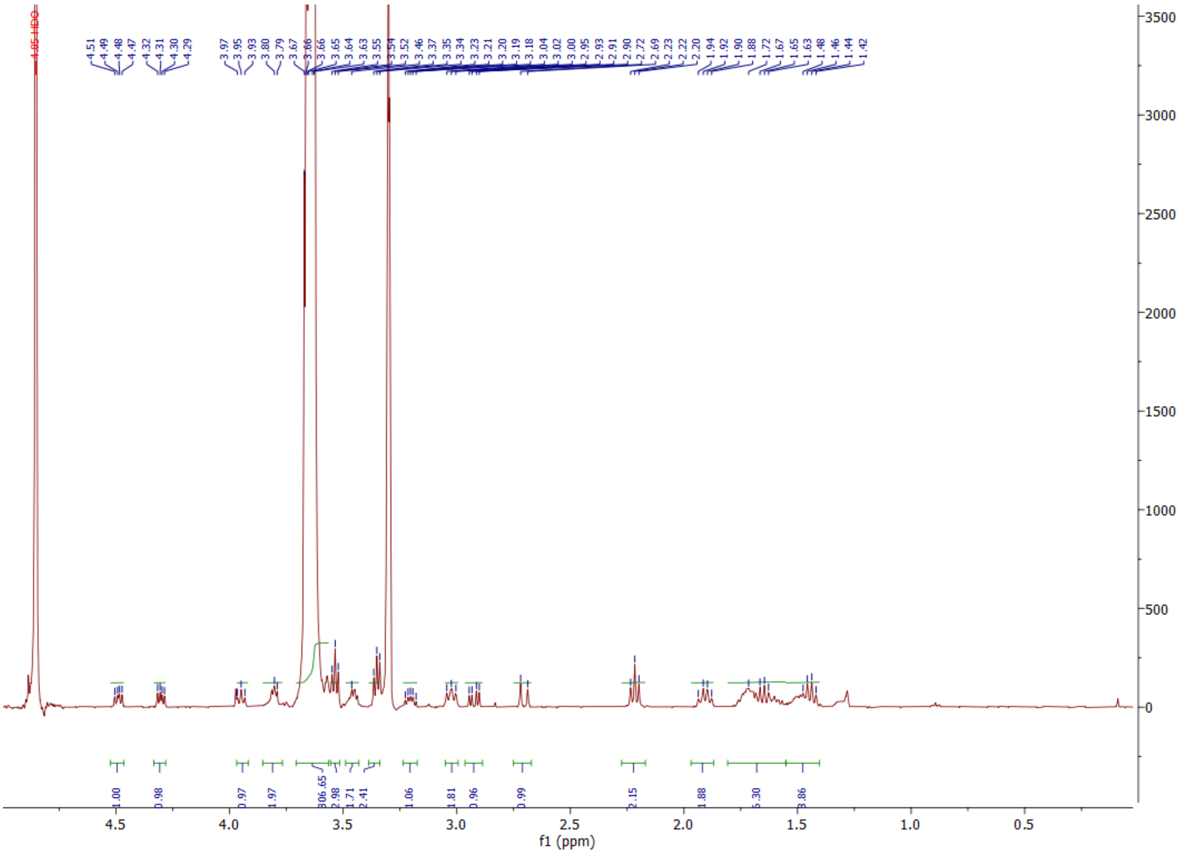


**Figure S4.** $^{1}$H-NMR of compound **4** in $CD_3OD$.

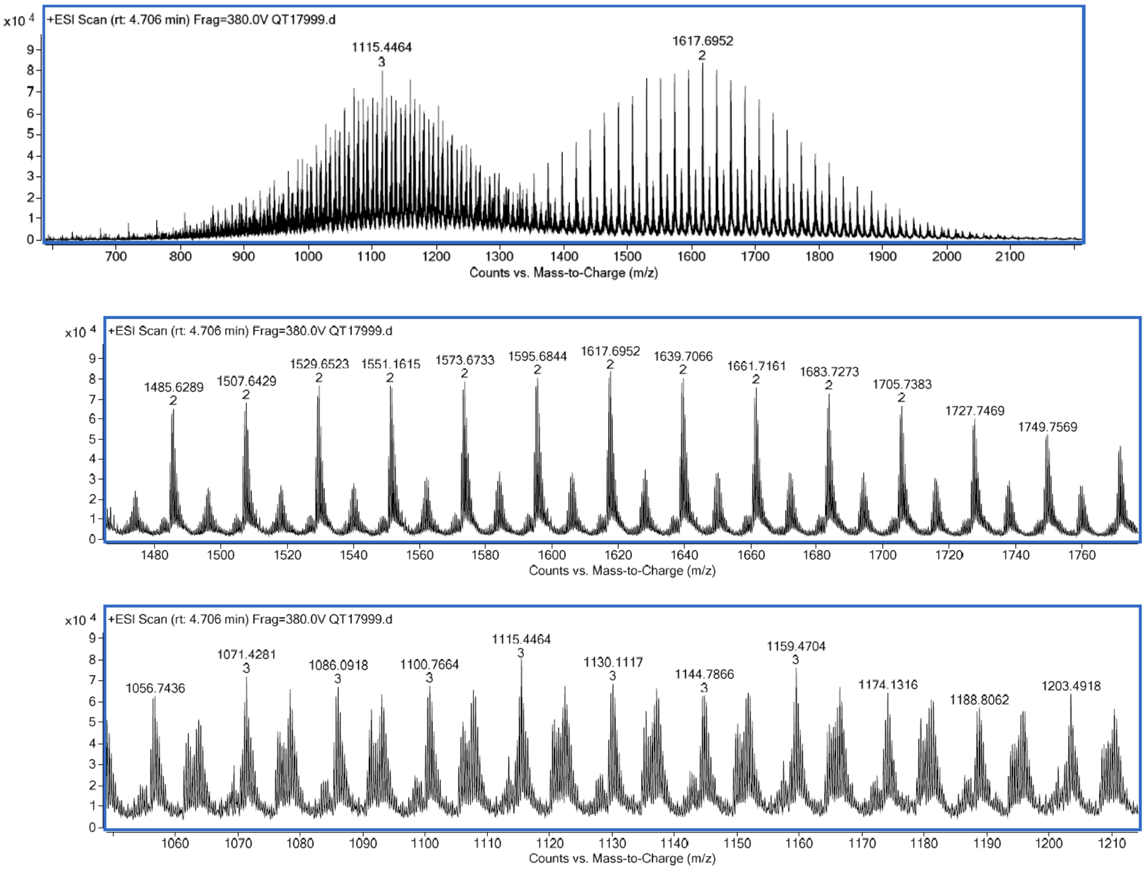


**Figure S5.** Mass spectra of compound **4** in $CD_3OD$.

**Formation non-functionalized nanoemulsions (NEs)**

NEs were produced by spontaneous nanoemulsification: 55 mg Vitamin E acetate containing 6% of NPC-Cholesterol with Cy5.5-TPB at 2% and 45 mg Kolliphor® ELP are mixed at 60 °C for 5 min. To the Oil:Surfactant mixture, 230 µL of filtered Milli-Q water is added, followed by 30 s of shaking with vortex, and 10 min with ThermoShaker (60°C, 1400 rpm).

**Nanoemulsion biotinylation**

55 mg Vitamin E acetate containing 6% of NPC-Cholesterol (with or without Cy5.5-TPB at 2%), Biotin-PEG-lysine (5 mg), and triethylamine (1.3 µL) were mixed in THF (500 µL), overnight at 60°C. THF and triethylamine were removed under reduced pressure. Kolliphor® ELP (45 mg) and filtered Milli-Q water (460 µL) were added according to the general protocol for nanoemulsion formulation. Then, 66 µL of biotinylated-NEs were diluted into 100 µL phosphate buffer (pH 8.5). Butylamine (1.6 µL) was added, followed by 2 hours of incubation on a ThermoShaker (room temperature, 400 rpm). Absorbance spectra were recorded before and after the addition of butylamine to determine the extent of the reaction with the biotin-linker. Finally, Biotin-NEs were purified by dialysis in Milli-Q water using a 14 kDa MWCO membrane.

**NPC-Cholesterol conversion**

NCP-Cholesterol (3.3 mg), Biotin-PEG-lysine (5 mg), and triethylamine (1.3 µL) were mixed in THF (500 µL), overnight at 60 °C. THF and triethylamine were removed under reduced pressure. The residue was dissolved in CDCl3 (100 µL) and transferred to a 3 mm diameter tube for $^{1}$H-NMR acquisition.

**Trastuzumab conjugation with biotin-NHS ester**

500 µL of Trastuzumab (32 µM in borate buffer) was mixed with 5 µL of NHS-biotin (100 mM in DMSO). The reaction was incubated overnight on a ThermoShaker (room temperature, 300 rpm). Afterward, purification was done by ultrafiltration (50 kDa MWCO) using PBS. The Biotin-Trastuzumab was stored at 4°C in DPBS, in the presence of 0.05% (w/v) of sodium azide, for conservation.

**Nanoemulsion functionalization with trastuzumab**

200 µL of NeutrAvidin (22 µM in DPBS) and 200 µL of biotinylated antibody (11 µM in DPBS) were incubated for 24 hours at room temperature in a DNA low-bidding Eppendorf. Then, 100 µL of biotinylated-NEs were added, followed by 24 hours of further incubation. Finally, antibody-NEs were purified by size exclusion chromatography using using Sephacryl S300 HR gel column, which is specially adapted for separating relatively large macromolecules or globular proteins in the range 10 to 150 KDa, and DPBS as an eluent.

**Biotin-NEs immobilization protocol**

8-well LabTek® chambers were treated with 500 µL KOH (1 M) each and incubated for 30 min. Similar treatments were done with 100 µL BSA-biotin (0.5 mg/mL) and 100 µL NeutrAvidin (0.5 mg/mL), each followed by 30 min incubation. Finally, the chambers were treated with biotinylated and non-biotinylated NEs as control, followed by 30 min incubation in darkness. Both biotinylated and non-biotinylated NEs were loaded with Cy5.5-TPB (2% with respect to the oil). All treatments were done at room temperature with DPBS washings in between.

**Cell experiments**

*Cell culture.* MDA-MB-231 and HCC-1954 were grown in RPMI1640 with L-Glutamine, supplemented with 10% fetal calf serum and penicillin (100UI/mL) and streptomycin (100µg/mL) at 37°C in a humidified atmosphere containing 5% $CO_2$.

*Cytotoxicity.* 2,500 MDA-MB-231 and 7,500 HCC-1954 were plated in a 96-well plate and let adhere overnight. The day after, medium was changed for medium containing increasing concentration of bare NEs or antibody-targeted NEs. After 72h, cells were washed, fixed with PFA 4% and stained for 1h with 2% Cristal violet. Excess of cristal violet was washed with tap water and plates were let dry before destaining by acetic acid. Cristal violet absorbance was measured at 595nm using a MultiSkan FC plate-reader.

*Nanoemulsion internalization.* 30,000 MDA-MB-231 or HCC-1954 were plated in an 8-well LabTek slide with coverlid and let adhere overnight. The day after, medium was changed for medium containing 1.1nM of bare NEs or antibody-targeted NEs in Opti-MEM. After 4h of incubation at 37°C, cells were wash and their membranes were stained with wheat germ agglutinin (WGA) Alexa-Fluor 488 (5 µg/mL in PBS) over 5 min at 37°C. After an additional wash, cells were fixed in PFA 4% and mounted in Fluoromount/DAPI. Slides were imaged using a 60X water-immersion objective on an inverted OIympus Spinning-disk. The comparions of SKBR3 vs MDA-MB 231 was done in another set of experiments. SKBR3 and MDA-MB 231 cells were grown in Dulbecco’s modified Eagle medium (DMEM, Gibco) supplemented with FBS (10%), sodium pyruvate (1 mM), L-glutamine (4 mM), and phenol red.

For imaging experiments, cells were seeded in 35 mm glass bottom dishes (Ibidi) and incubated over 2 days, at 37 °C under a humidified 5% CO2 atmosphere. Then, cells were washed with DPBS, followed by 30 min incubation in darkness with Hoechst 33342. After washing with DPBS, cells were incubated with Antibody-NEs (0.2 nM in OptiMem) on ice for 1 hour. Finally, they were washed with OptiMem.

**Statistical analysis**

Statistical analysis was performed with GraphPad Prism 9 software. The normal distribution of the data was tested using the Shapiro–Wilk normality test. When comparing two groups, a Mann-Whitney analysis were used, for more than 2 groups a Kruskal-Wallis test followed by the original FDR method of Benjamini and Hochberg post-test was used. $p<0.05$, *; $p<0.01$, ** . $p<0.001$, ***; $p<0.0001$, ****. Data in graphs are presented as mean ± Standard Deviation.

### Supplementary Figure legends

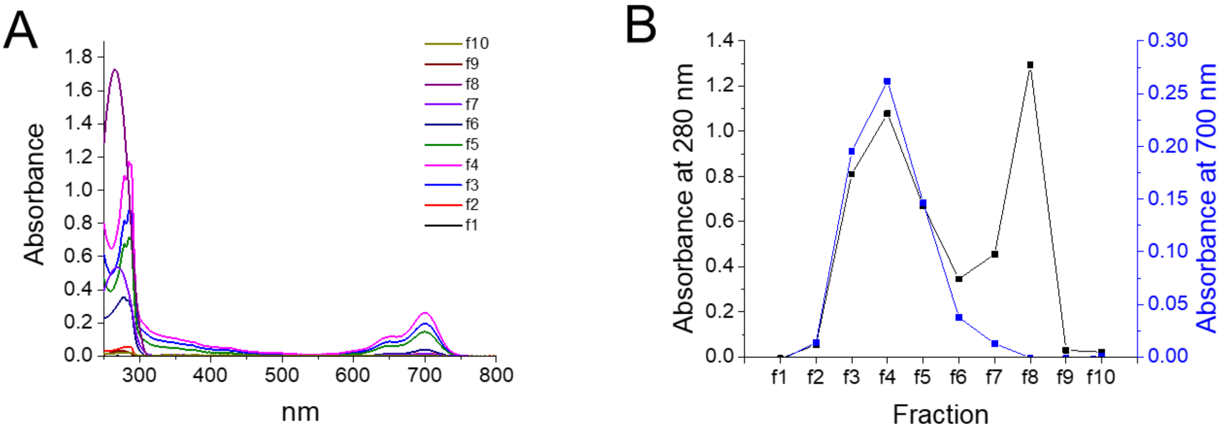


**Figure S6.** (A) Absorption spectra of different fractions of non-purified mixture containing NEs, neutravidin and biotinylated mAbs. (B) Absorbance value for different fractions recorded at 280 and 700 nm.

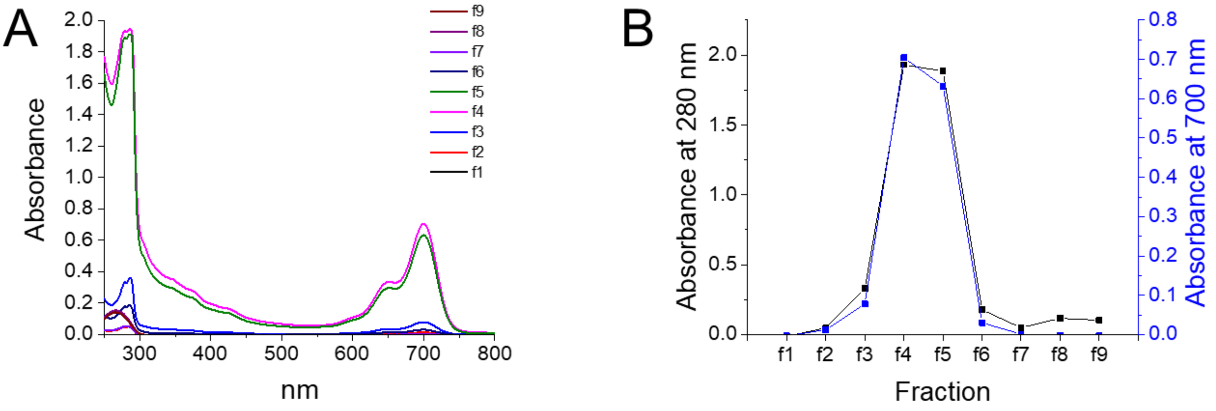


**Figure S7.** (A) Absorption spectra of different fractions of NEs conjugated with mAbs after purification. (B) Absorbance value for different fractions recorded at 280 and 700 nm.

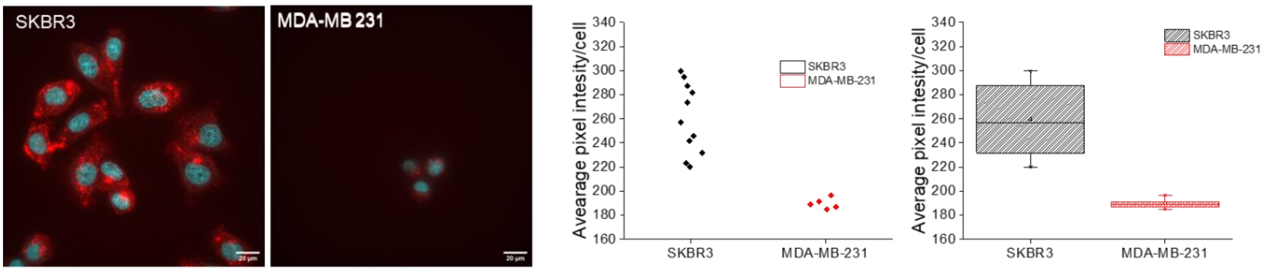


**Figure S8.** Epifluorescence microscopy images from SKBR3 and MDA-MB 231 cells after 1h incubation on ice with Antibody-NEs (0.2 nM) containing 2% of Cy5.5-TPB. Data on the left are the quantitative image analysis.

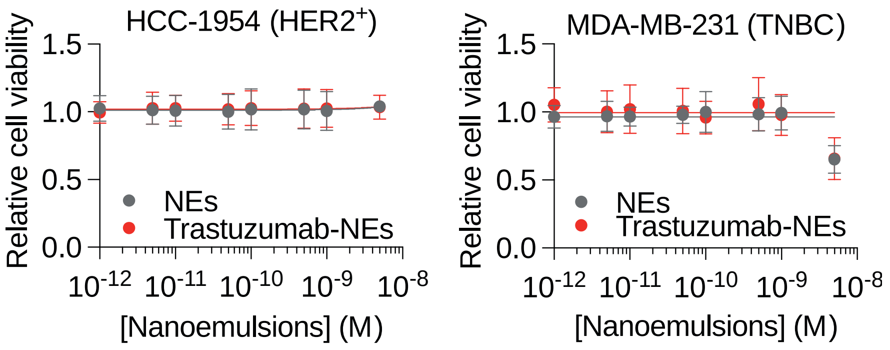


**Figure S9.** Nanoemulsions cytotoxicity was measured after a 72h treatment with NEs increasing concentrations of HER2-amplified HCC-1954 (left) and HER2$^{low}$ MDA-MB-231 (right) cells. Cytotoxicity was assessed using a crystal violet staining.

**Table S1.** Size of NEs measured by DLS.

| NEs type | Diameter (nm) | Polydispersity index (PdI) |
|---|---|---|
| Non-functionalized (6 wt. % NPC-Chol and 2 wt. % Cy5.5-TPB) | 53.3 ± 0.2 | 0.13 ± 0.02 |
| Biotinylated (no dye) | 64 ± 4 | 0.20 ± 0.03 |
| Biotinylated (2 wt. % Cy5.5-TPB) | 53 ± 2 | 0.16 ± 0.01 |
| Antibody (2 wt. % Cy5.5-TPB) | 56 ± 2 | 0.12 ± 0.03 |
